# Synthetic yeast genome SCRaMbLEing uncovers a new role for ribosomal proteins in genetic code expansion

**DOI:** 10.1101/2025.08.31.673356

**Authors:** James E. J. Sanders, Stefan A. Hoffmann, Ewan R. Moody, Mark McCullough, Anthony P. Green, Yizhi Cai

## Abstract

The translation of proteins with non-canonical amino acids (ncAAs) has emerged as a powerful technology for embedding new functional elements into proteins, enabling the development of novel enzymes, materials, and biopharmaceuticals. However, the utility of this approach has been hindered by weak translation efficiencies. To address this challenge, we sought to substantially improve orthogonal translation in *Saccharomyces cerevisiae*. We first evaluated recently described ΔNPylRS-class pyrrolysyl-tRNA synthetase systems and identified a homolog with ∼5.4-fold higher activity than the best previously reported pyrrolysyl system. Building on this advance, we leveraged the SCRaMbLE system in the semi-synthetic yeast strain Syn6.5 to generate structural genomic variation, and identified strains with enhanced ncAA incorporation. Pooling genomic alterations across enhanced strains, we identified an association for the deletion of several ribosomal protein genes with the increased production of ncAA-containing protein. These findings demonstrate a previously unrecognized role for ribosomal proteins in enabling alternate genetic codes.

## 1 Introduction

Genetic code expansion enables the translation of proteins equipped with moieties not found within the standard set of 20 canonical amino acids (cAAs)^1^. Translation of these non-canonical amino acids (ncAAs) is coordinated by orthogonal aminoacyl-tRNA synthetase-tRNA pairs that enable site-specific translation, often directed towards the amber UAG stop codon. Expanding upon the building blocks of translation has opened up new possibilities for functional protein design, leading to the development of new biotherapeutics^2^, biomaterials^3^, biocatalysts^4^, and biocontainment strategies^5^.

Despite great progress, transition of genetic code expansion research into large scale applications is limited by the relatively poor translation efficiencies of ncAAs compared to cAAs. Inclusion of even a single ncAA within a protein can lead to substantial reductions in protein yield^6^. These issues result in high production costs of modified proteins that become even more problematic when incorporating multiple ncAAs into proteins as required for some applications. The incompatibility of ncAAs within existing translational networks is likely because the evolution of translation has been tightly coupled to the canonical set of amino acids. Indeed, translation is one of the oldest and most conserved functions within the cell, dating back ∼4.2 billion years to the last universal common ancestor from which all extant life has evolved^7^. In this time, it has evolved to excel at translating the cAAs while preventing mistranslation. In this respect, genetic code expansion challenges the endogenous machinery and regulatory systems that have evolved to optimize and safeguard canonical translation.

In *Escherichia coli*, efforts to enhance the incorporation of ncAAs have been aided by advances in synthetic genomics. For example, the removal of all endogenous TAG stop codons from synthetic *E. coli* genomes has enabled the deletion of the release factor, RF1, reducing competition at the site for orthogonal translation systems^8,9^. However, the monopolization of codons alone does not facilitate efficient translation of ncAAs. Translation is a complex orchestration of hundreds of proteins and RNAs that collectively facilitate the rapid, high-fidelity translation of cAAs. To date, research in improving ncAA incorporation has focused largely on the tRNA-aminoacyl-tRNA synthetase pair^10^. Although this strategy has made great progress in improving ncAA incorporation, achieving truly efficient translation of ncAAs will likely require that the intricate network of interactions governing canonical translation be holistically adapted towards alternate amino acid moieties.

Exploring combinatorial perturbations to this network and their impacts on translation is a challenging task. In *Saccharomyces cerevisiae*, screening single gene deletions for improved ncAA translation has yielded some success, although candidate genes were cryptic in their involvement in ncAA translation^11^. To progress in guiding this network towards accepting ncAAs, directed evolution techniques that capture the complexity of epistatic interactions and finely-tuned gene dosage compensations within this cellular process will have to be employed. Recently, within the international synthetic yeast (Sc2.0) project, synthetic versions of all 16 chromosomes of *S. cerevisiae* have been assembled^12,13,14^. Bottom-up genome synthesis has enabled the incorporation of new-to-nature design features. This includes the replacement of all TAG stop codons to the synonymous TAA triplet and the integration of over 4,000 symmetrical loxP (loxPSym) sites downstream of every non-essential gene. By positioning these sites throughout the genome, inducible Cre recombinase activity can generate stochastic genome-wide deletions, duplications, inversions, and translocations of chromosomal segments through a function called Synthetic Chromosome Recombination and Modification by LoxP-Mediated Evolution (SCRaMbLE).

Here, we explored whether SCRaMbLE can effectively re-wire the yeast interactome to efficiently translate new amino acid moieties. We first expanded upon the tools for genetic code expansion in *S. cerevisiae* by isolating seven new pyrrolysyl systems with novel ΔNPylRS domains that function in yeast. The most active of these variants, from the *Methanomassiliicoccales archaeon* isolate RumEn M1, we found to be ∼5.4-fold more active for the incorporation of N*ɛ*-Boc-L-Lysine (BocK) than the current best-in-class system from *Methanomethylophilus alvus*^15^. We then improved upon the activity of this system in yeast by SCRaMbLEing the synthetic yeast strain Syn6.5^13^. By developing a high-throughput evolution pipeline utilizing a novel *URA3* auxotrophy reporter assay, we showed that SCRaMbLE can improve the suppression of two sites by *>*250% compared to the parental unSCRaMbLEd strain. Finally, we pooled the genomic information across 14 evolved strains and isolated several new genomic determinants for genetic code expansion that promote the translation of ncAAs in wild-type strains. In particular, we found new roles for ribosomal proteins and translationally-adjacent genes in modulating the production of ncAA-containing proteins.

## 2 Results

### 2.1 An Expanded Pyrrolysyl Toolbox for *Saccharomyces cerevisiae*

A growth-based reporter was developed to assess the activity of the orthogonal translation systems (*Fig. 1a*). Here, two sites within the protein orotidine 5’-phosphate decarboxylase (Ura3p), essential for uracil biosynthesis, were identified to be permissible for amino acid substitution (*Fig.* S1). Codons of protein positions E143 and K209 were replaced with the amber stop TAG triplet, creating *URA3-E143*-K209\** (*URA3-2TAG*). The rationale of this assay is that highly active orthogonal translation systems (OTSs) co-transformed with this assay are able to suppress both amber stop sites with the supplied ncAA leading to full-length protein production, supporting growth of *ura3*Δ strains on uracil-deficient media.

Eleven ΔNPylRS-class pyrrolysyl homologs, previously characterized in *Escherichia coli* ^16^, were tested with cognate tRNA in the wild-type *S. cerevisiae* strain, BY4742. The ncAA N*ɛ*-Boc-Llysine (BocK), a natural substrate for pyrrolysyl synthetases, was used to assess these systems. As shown in *Figure 1c*, eight out of the eleven displayed some degree of activity with the *URA3* assay when grown in the absence of uracil and in the presence of 1 mM BocK. These active homologs were found across class A and B ΔNPylRS systems (*Fig. 1b*). Growth on uracil-deficient media without BocK would indicate promiscuous activity. None of the eight active pairs supported growth without uracil and BocK. Overall spot growth and colony size on uracil drop-out media supplemented with ncAA suggest the pyrrolysyl system from the *Methanomassiliicoccales archaeon* isolate RumEn M1 (RumEn) to be the most active among this collection for the incorporation of BocK.

A fluorescence-based assay^17^ was used to quantify the activity of the homologs for the ncAA-dependent production of GFP. As shown in *Figure 1d*, the RumEn OTS was also the most active variant for the production of GFP, followed by the pyrrolysyl system from *Candidatus Methanomethylophilus* sp. 1R26 (1R26), closely matching the growth-assay results. It was noted that RumEn exhibited a ∼5.4-fold improvement over the activity of the previous best-in-class pyrrolysyl system from *Methanomethylophilus alvus* (*Alv*)^15^.

**Figure 1:**
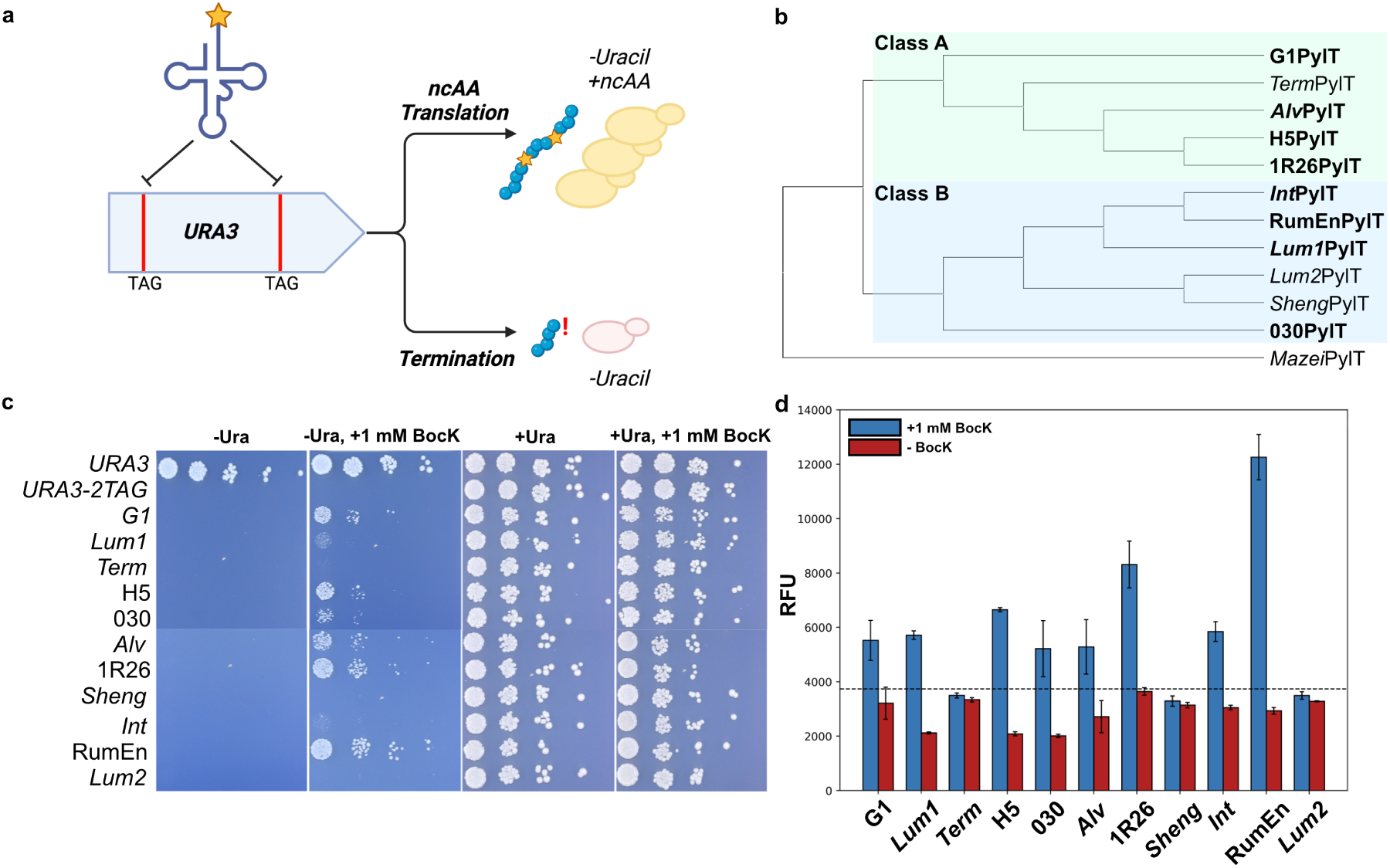
Characterization of **Δ**N-Class Pyrrolysyl Homologs. **a** Protein positions E143 and K209 in the coding sequence of the *URA3* gene were replaced with TAG stop codons, creating *URA3-2TAG*. When grown on uracil-deficient media, presence of the active orthogonal translation system and a suitable non-canonical amino acid allows the strain to grow. Absence of either of these factors leads to lack of growth. **b** Maximum likelihood tree depicting the two classes of tRNA among the pyrrolysyl homologs with active systems in bold. The tRNA of the +NPylRS orthogonal translation system from *Methanococcus mazei* was used as an out-group. **c** Strains of *S. cerevisiae* BY4742 (MAT*α his3*Δ*1 leu2*Δ*0 lys2*Δ*0 ura3*Δ*0*) were co-transformed with the *URA3-2TAG* growth assay and an orthogonal translation system. As a control, BY4742 was transformed with a corresponding empty backbone (pRS425) in combination with either the TAG-less *URA3* or *URA3-2TAG* plasmid. Strains were spotted on SC-His-Leu in the presence or absence of BocK/uracil and incubated for three days. **d** Pyrrolysyl homologs were tested in BY4742 with the pRS413-BXG assay^17^ for incorporation of BocK. Strains were grown without or with 1 mM BocK. The dotted line represents the threshold for activity above auto-fluorescence of a control strain transformed with two empty plasmids. Shown are mean sfGFP fluorescence values ± SD (n=3).

### 2.2 SCRaMbLE Improves BocK Incorporation in Syn6.5

Having identified a highly active pyrrolysyl system, RumEnPylRS-RumEnPylT, we next sought to optimize the genetic background of *S. cerevisiae* to improve the translation of BocK with this OTS by generating genomic diversity through SCRaMbLE. Here, the addition of *β*-estradiol promotes the nuclear import of a Cre recombinase tagged with an estradiol binding domain^18^. Once imported, Cre is able to act upon the loxPSym sites embedded across the synthetic chromosomes to create deletions, inversions, duplications, and translocations of loxPSym segments^19^. To effectively isolate yeast strains with structural genomic variations that support enhanced ncAA incorporation, a high-throughput lab automation pipeline was developed to quickly filter and identify top-performing mutants from large SCRaMbLEd pools. This pipeline was based on selection with the *URA3* growth assay described previously to enable progressive filtering of strains in a three-stage cycle depicted in *Figure 2*.

Directed genome evolution of Syn6.5 carrying the synthetic chromosomes II, III, V, VI, IXR, X, and XII with the SCRaMbLE pipeline enabled considerable improvements in ncAA incorporation after one to two rounds of mutagenesis. As shown in *Figure 3a*, growth on selective -Ura, + 1 mM BocK media improved considerably for SCRaMbLEd strains indicating an increase in the production of Ura3p containing the ncAA. To quantify this improvement, strains were assessed by growth rate in liquid culture. Growth rates in selective conditions were found to be up to ∼250% that of parental Syn6.5 (*Fig. 3b*). For many strains, it was found the increased ability to translate ncAA-containing proteins was accompanied by a slight growth defect under non-auxotrophic conditions (*Fig. 3c*).

### 2.3 Deconvolution of SCRaMbLEd Genomes

SCRaMbLEd genomes are inherently difficult to de-tangle mechanistically^19,20^. This is because of i) the high number of passenger mutations often associated with structural re-arrangements, ii) the ambiguous effect of some structural re-arrangements on gene dosage, iii) the accumulation of multiple SCRaMbLE events in any one strain, and iv) the potential for phenotypes deriving from epistatic interactions. To overcome these challenges, we mapped the changes to the yeast protein-protein interactome within a pool of SCRaMbLEd strains to deduce genetic interactions underpinning the phenotypes. Here, cellular functions that show a high clustering of SCRaMbLE mutations may be highlighted, allowing for focused efforts in screening for driver genetic determinants from SCRaMbLEd genes within these processes.

**Figure 2:**
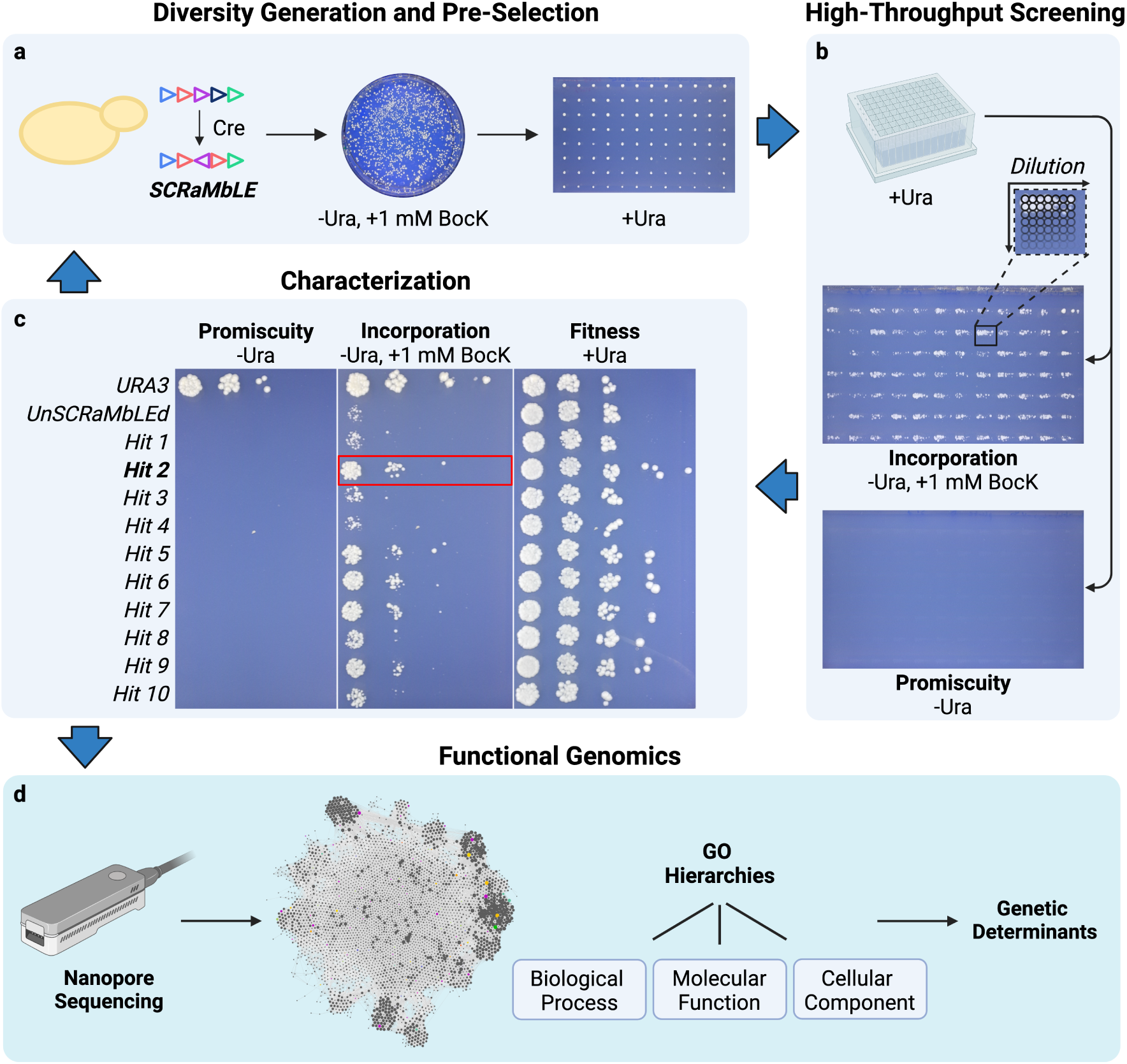
SCRaMbLE Evolution Pipeline. Pipeline to identify strains that exhibit improved ncAA incorporation. **a** SCRaMbLEd cultures were plated on selective -Ura, +1 mM BocK media and incubated until colony formation. Large colonies were picked with an automated colony picker into 96-array format on non-selective media. **b** Pre-screened colonies were then grown to saturation and spotted on -Ura, +1 mM BocK and -Ura media in 7x7 grids to assess for incorporation activity and promiscuity at high throughput. **c** Strains with large 7x7 grid patches on -Ura, +1 mM BocK were cultured to saturation and spotted onto the four combinations of Ura/BocK to assess strain characteristics. Top-performing strains from this characterization could then be re-entered into the pipeline for continued SCRaMbLE evolution or **d** submitted for analysis by whole-genome sequencing to deconvolute SCRaMbLEd genotypes.

**Figure 3:**
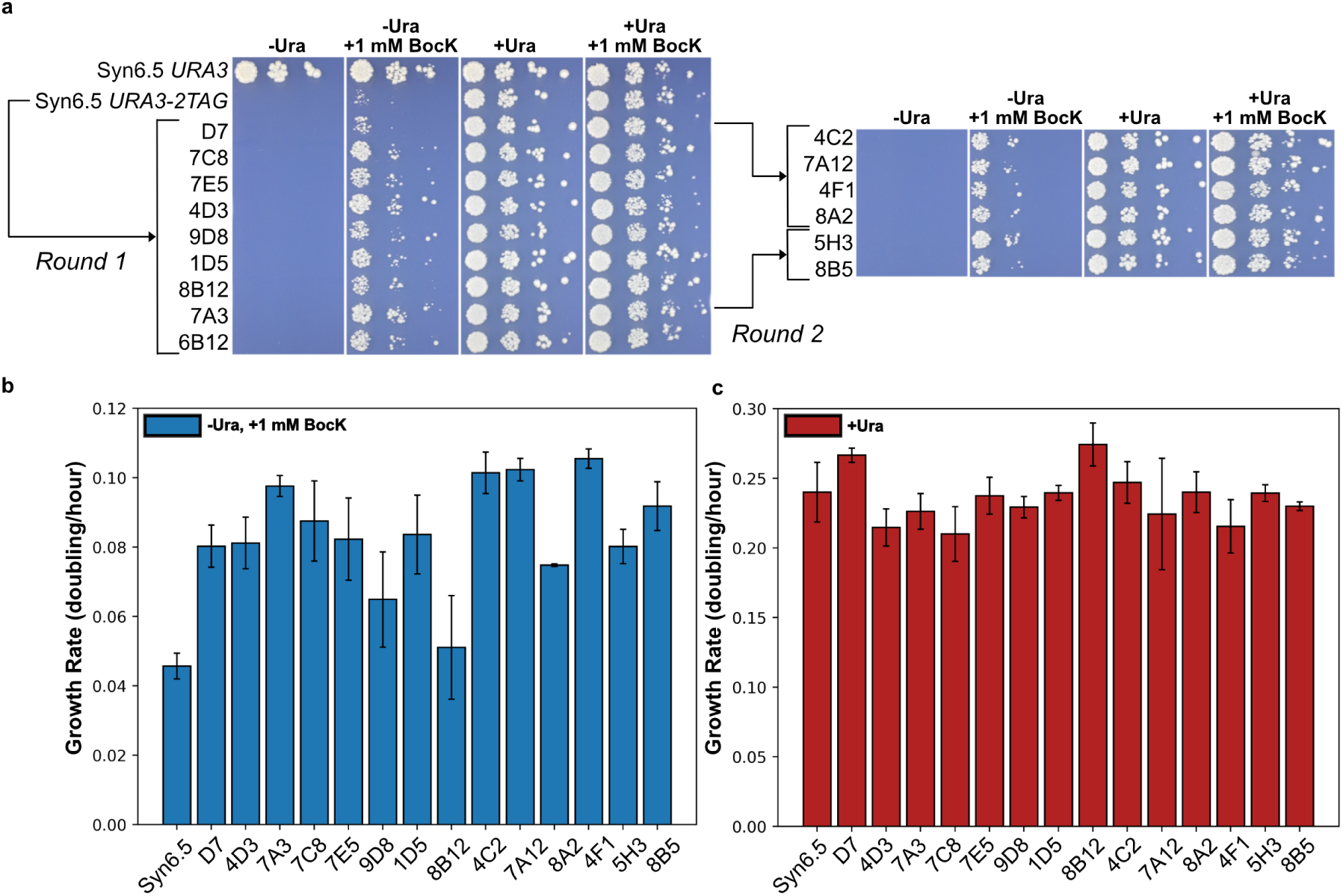
Characterization of SCRaMbLEd Strains. **a** SCRaMbLEd strains co-transformed with the *URA3-2TAG* growth assay and RumEn orthogonal translation system were spotted on SC-His-Lys with and without uracil or 1 mM BocK. Alongside evolved strains, parental Syn6.5 was spotted as a baseline control. **b** Growth rates of round 1 (D7-8B12) and round 2 (4C2-8B5) SCRaMbLEd strains transformed with *URA3-2TAG* assay in selective SC-His-Lys-Ura +1 mM BocK media. **c** Growth rates of SCRaMbLEd strains in non-auxotrophic SC-His-Lys (+Ura) media. Shown are mean rates ± SD (n=3).

Fourteen strains with improvements to ncAA-containing protein production were sequenced using Nanopore long-read sequencing to determine underlying SCRaMbLE events (*Fig. 4c*). All the genes affected by SCRaMbLE events were pooled and mapped onto the protein-protein interaction network shown in *Figure 4a*. Here, every node represents a protein with an edge connecting it to an interacting partner. From the graph, it can be seen that major cellular functions form clusters with high degrees of inter-protein interaction.

**Figure 4:**
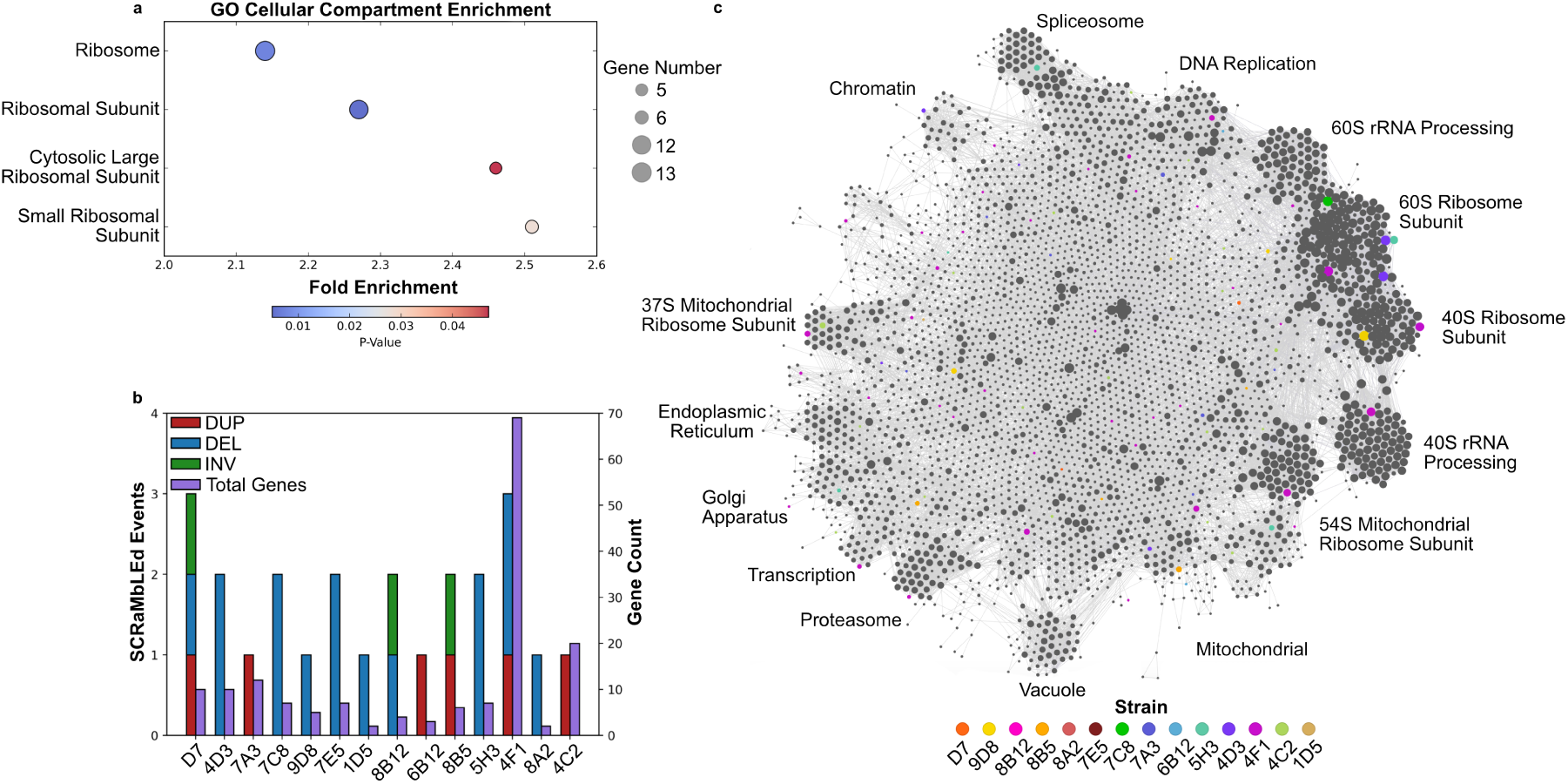
Gene-Set Enrichment Analysis of SCRaMbLEd Strains. **a** SCRaMbLEd genes were assessed for statistical-enrichment by the cellular compartment gene ontology (GO)-Term. Enrichment was assessed relative to a reference list of genes found on synthetic chromosomes. Terms with a *>*2-fold enrichment associated with P-values *<*0.05 were plotted. **b** Breakdown of the number of duplication (DUP), deletion (DEL), and inversion (INV) SCRaMbLE events per strain with the number of genes affected by SVs across all SCRaMbLE events. **c** SCRaMbLEd events mapped onto yeast protein-protein interactome with each node in the graph representing a protein with an edge symbolizing the physical interaction with another protein. SCRaMbLEd protein genes were color-coded by strain.

We hypothesized that cellular processes affected by SCRaMbLE events across multiple strains may indicate a causative connection to the production of proteins containing ncAAs. Within the interactome of the pooled SCRaMbLEd strains, both subunits of the cytosolic ribosome were shown to have high clustering of SCRaMbLE events across multiple strains. These networks are predominantly composed of ribosomal proteins (RPs) which bind with rRNA to aid in structure and function^21^. In *S. cerevisiae*, the small 40S and large 60S subunits are formed with 33 and 46 RPs respectively^22^. The high number of SCRaMbLE events affecting RP genes was found to be of particular interest given the central role of the ribosome in translation. To support the qualitative assessment of the interactome, gene set enrichment analysis was performed using the gene-ontology (GO) framework. As shown in *Figure 4b*, in the GO cellular component domain significant enrichment was found for both ribosomal small and large subunits.

### 2.4 Deletion of Ribosomal Protein Genes and *TMA22* Increase Production of BocK-Containing Protein

To assess the influence of ribosomal proteins on the incorporation of BocK, ribosomal protein genes associated with SCRaMbLE events in selected strains were characterized as single-gene knockouts in the haploid, wild-type background BY4742. We opted to reverse-engineer SCRaMbLE improvements in the fitter wild-type background with the ultimate goal of creating an improved production chassis for proteins containing ncAAs. To increase assay stringency to account for the increase in basal fitness, strains were characterized using a *URA3-3TAG* assay (*URA3-E71*-E143*-K209\**) requiring the incorporation of three ncAAs to produce full-length Ura3p.

As shown in *Figure 5a* and *Figure 5e*, it was found that single knockouts of *RPS4A*, *RPS6B* and *RPL19A* encoding small ribosomal subunit proteins eS4, eS6 and large ribosomal subunit protein eL19, supported efficient incorporation of three BocKs into Ura3p. These three genes were found to be lost within separate SCRaMbLEd strains (9D8, 4F1, and 4D3, respectively) as a result of Cre-based deletions of loxP segments. Interestingly, the ribosomal protein knockout, which most improved BocK translation in BY4742, *rps6b*Δ, was derived from one of the best performing SCRaMbLEd variants (4F1).

Due to an evolutionary conserved duplication event, the majority of RPs are encoded by two genetic paralogs, denoted as the A and B isoforms^23^ which show high sequence identity. Despite sharing similar or identical sequences, several reports have pointed to paralog-specific effects. For example, localization of the transcription repressor, Ash1, to yeast daughter cells was found to require a specific set of paralogs^24^. We did not find paralog-specific effects for the deletion of eS4 or eS6 genes, with the deletion of either paralog contributing to increased Ura3-3BocK production (*Fig. 5d*). However, for eL19, it was found that only deletion of *RPL19A* led to phenotypic improvement, while deletion of *RPL19B* resulted in a stronger reduction in fitness in both auxotrophic and non-auxotrophic conditions.

**Figure 5:**
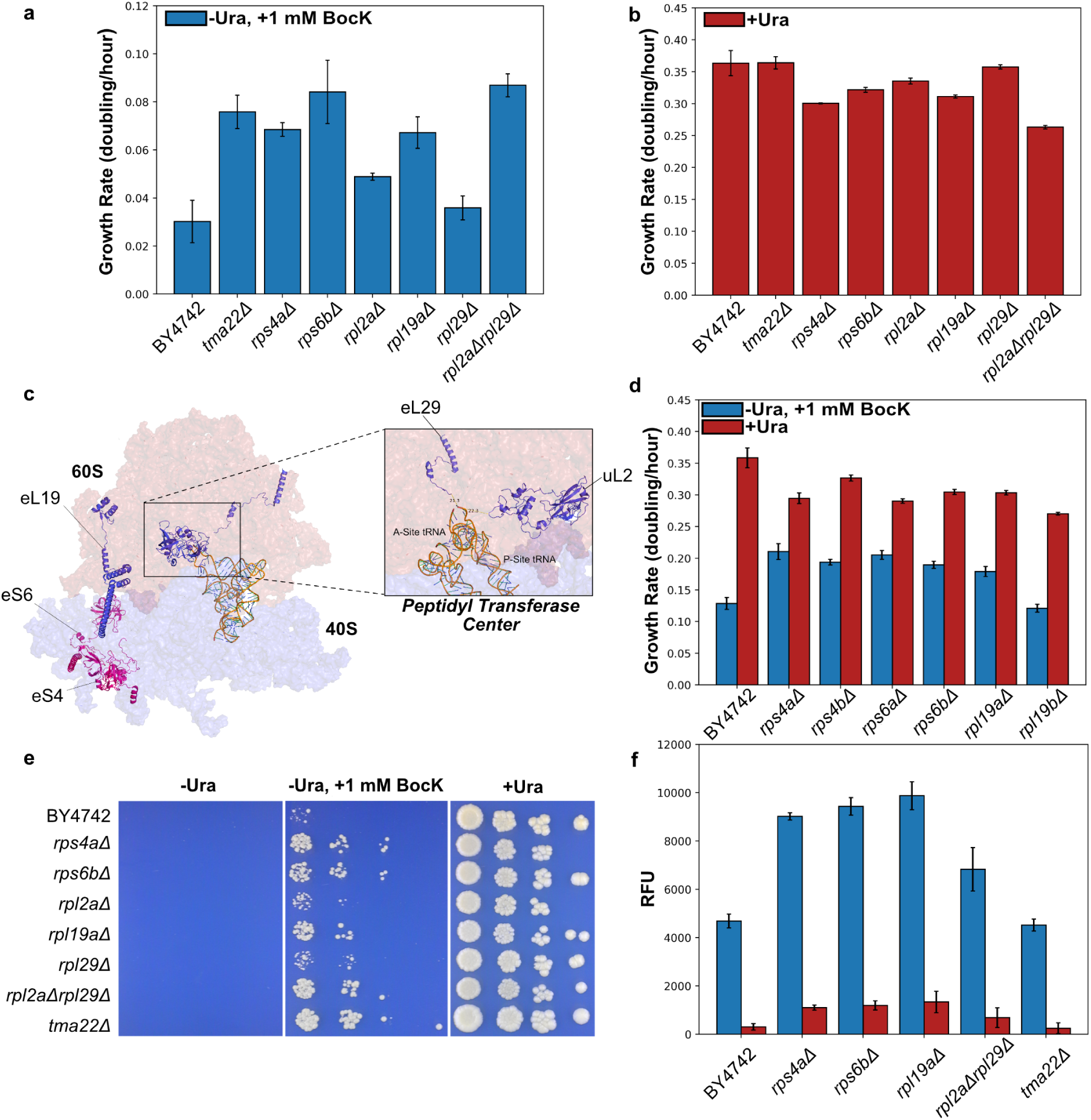
Effect of Ribosomal Protein Deletion on BocK Translation. **a** Growth rates of BY4742 and single-gene deletion strains with the *URA3-3TAG* assay in selective SC-His-Lys-Ura +1 mM BocK medium. Shown are mean rates ± SD (n=3). **b** Growth rates of BY4742 and single-gene deletion strains in non-auxotrophic SC-His-Lys media. **c** Location of SCRaMbLEd ribosomal proteins on the ribosome (PDB: 6TNU^25^). **d** Growth rates of ribosomal protein paralog deletion strains in SC-His-Lys media without uracil, without uracil with 1 mM BocK, and with uracil while expressing the *URA3-2TAG* reporter. **e** Spot assay of BY4742 and single-deletion strains on SC-His-Lys-Ura, SC-His-Lys-Ura +1 mM BocK and SC-His-Lys media. **f** End-point fluorescence of strains transformed with pRS423-msGFP-N149TAG and pRS425-RumEnPylT-RumEnPylRS grown with and without 1 mM BocK. Shown are mean relative fluorescence units (RFU) ± SD (n=4).

For two RPs, uL2 and eL29 encoded by *RPL2A* and *RPL29*, single knockouts of either gene led to either modest or no improvements in BocK-dependent growth. However, in SCRaMbLEd strains these genes were only found removed together in a single deletion event on chromosome VI. This particular event was mirrored across three of the sequenced strains (5H3, 7E5, and 7C8), suggesting a strong enrichment for this genome re-arrangement among the SCRaMbLE population and a causative role of the double knockout for increased ncAA translation. Upon inspection of the two proteins’ location within the ribosome, it was found that uL2 and eL29 are in close proximity to each other near the peptidyl transferase center (*Fig. 5c*). We hypothesized that knocking out both proteins may create a synergistic effect that aids BocK translation. Indeed, deletion of both *RPL2A* and *RPL29* led to super-additive improvements in the production of Ura3p-3BocK, increasing BocK-dependent growth rate by ∼300% compared to BY4742.

In addition to the deletion of genes encoding ribosomal proteins, we also found that knockout of the Translational-Associated Machine 22 (*TMA22*) gene led to substantial improvements in the production of Ura3-3BocK. As shown in *Figure 5a*, *tma*22Δ led to similar improvements as that of top-performing RP gene deletions but with no observable fitness defects when grown under non-auxotrophic conditions (*Fig. 5b*).

To validate the results of the growth-assay, we went on to characterize these strains with a GFP fluorescence reporter. Due to the fitness defect imparted by ribosomal protein deletion, we developed an alternative assay to that used to characterize the homologs (*Fig. 1d*) in which we found the switch to galactose induction exacerbated the existing fitness defect of the deletion strains. Here, we expressed msGFP^26^ under control of the constitutively active TDH3 promoter on a high-copy plasmid backbone with ncAA-incorporation at protein site 149. As shown in *Figure 5f*, an increase in GFP production was found for all of the RP gene deletions compared to parental BY4742. This amounted to a ∼200% improvement for the *rps4a*Δ, *rps6b*Δ and *rpl19a*Δ strains and a ∼145% improvement for the *rpl2a*Δ*rpl29*Δ strain. We did note that for many of these deletion strains a slight increase in msGFP production in the absence of BocK was observed, which indicates a potential increase in non-specific readthrough. For the *tma22*Δ strain, we did not observe any increase in msGFP relative to the wild-type background, suggesting that the enhancing effects of this deletion are assay-dependent. To confirm the presence of BocK within msGFP, we purified the protein from wild-type and knockout strains for analysis with mass spectrometry. As presented in *Fig. S2*, all strains expressing msGFP-N149TAG showed a protein mass spectrum corresponding to the successful incorporation of BocK.

**Figure 6:**
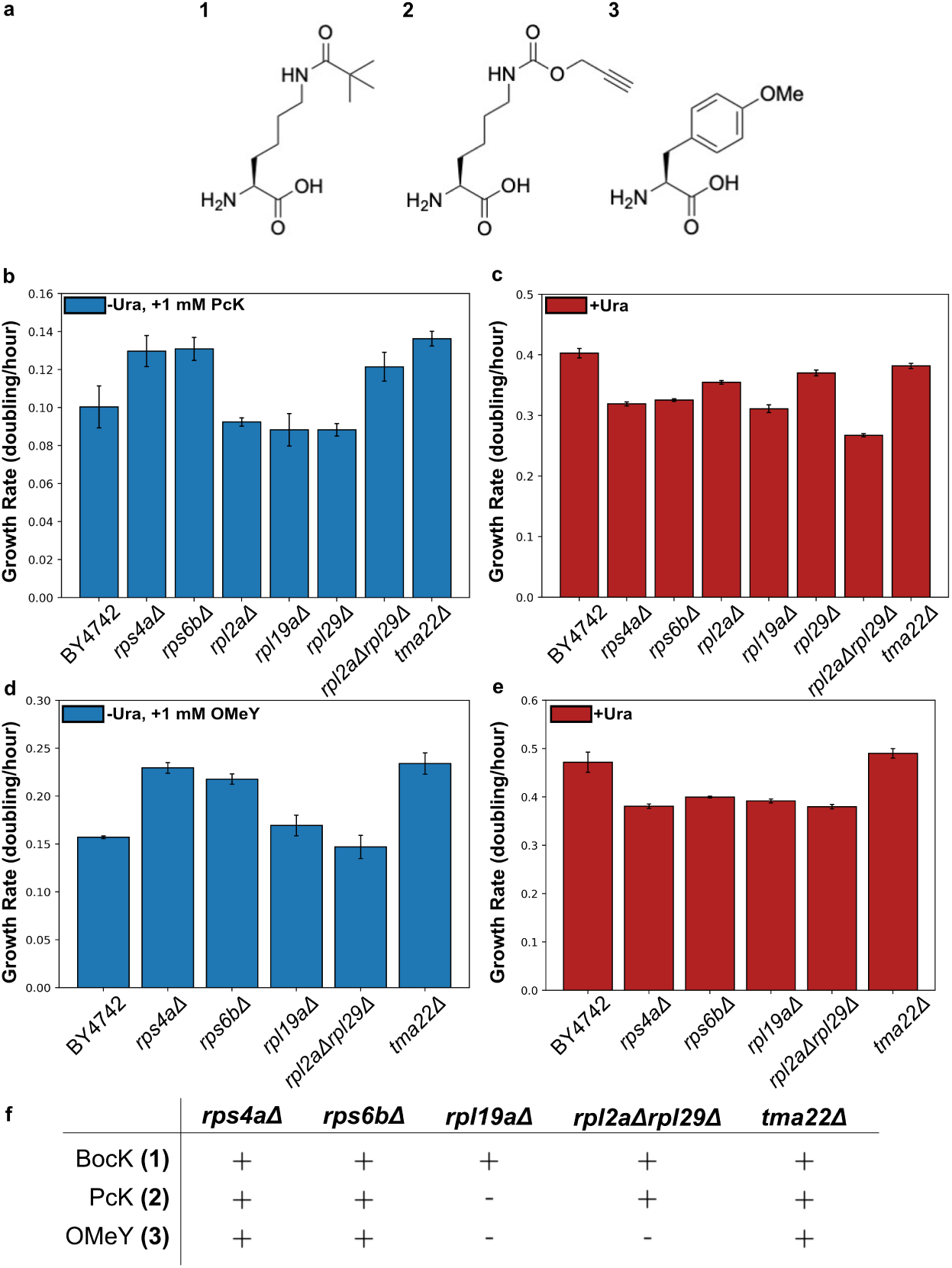
Effect of Genetic Determinants across ncAAs and Translation Systems. **a** Structure of ncAAs used in study: **1** N*ɛ*-Boc-L-lysine (BocK), **2** N^6^-((prop-2-yn-1-yloxy)carbonyl)-L-lysine (PcK), **3** O-methyl-L-tyrosine (OMeY). **b** Growth rates of BY4742 and deletion strains transformed with the RumEnPyl OTS and *URA3-3TAG* assay in selective SC-His-Lys-Ura +1 mM PcK and **c** non-auxotrophic SC-His-Lys medium. **d** Growth rates of BY4742 and deletion strains transformed with the *Ec*OMeY OTS and *URA3-3TAG* assay in selective SC-His-Leu-Ura +1 mM OMeY and **e** non-auxotrophic SC-His-Leu medium. Rates shown as mean ± SD (n=3). **f** Table of genetic determinant effectiveness across ncAA, (+) being effective, (-) being non-effective.

### 2.5 Genetic Determinants can be ncAA and OTS Specific

To assess the sensitivity of genetic determinants to different ncAAs and orthogonal translation components, knockout strains displaying improved BocK incorporation were further evaluated for their ability to encode with N^6^-((prop-2-yn-1-yloxy)carbonyl)-L-lysine (PcK) and O-methyl-L-tyrosine (OMeY). PcK contains an alkyne functional handle for bio-orthogonal conjugations^27^ and is a close analog of BocK that can be encoded into proteins using wild-type pyrrolysyl OTSs. In this case, we found the majority of deletions tested led to improved PcK incorporation over parental BY4742, with the exception of *rpl19a*Δ (*Fig. 6b*).

To investigate whether the isolated genetic determinants would aid in the translation of a chemically dissimilar ncAA incorporated by an alternative OTS, strains were assessed for activity with the *Ec*OMeY system and OMeY^28^. As shown in *Figure 6d*, all determinants except *rpl19a*Δ and *rpl2a*Δ*rpl29* Δ supported improved translation for OMeY over parental BY4742. As summarized in *Figure 6f*, we demonstrate that genetic determinants for genetic code expansion are not necessarily universal across ncAA and OTS. However, while *rpl19a*Δ and *rpl2a*Δ*rpl29* Δ appeared to be sensitive to ncAA and OTS, it was found that *rps4a*Δ, *rps6b*Δ, and *tma22* Δ provided broad support for the improved incorporation of ncAAs into Ura3p.

### 2.6 Discussion

Enhancing the capabilities of strains to incorporate new amino acids efficiently will enable continued advancement in the design and economical production of ncAA-containing proteins. However, engineering strains to efficiently translate ncAAs requires advancements in our understanding of genetic code expansion and translation. In this research, we expanded the toolbox of pyrrolysyl orthogonal translation systems in yeast, identifying seven new ΔPylRS-tRNA pairs that function in *S. cerevisiae*. From this set, we identified the OTS from the *M. archaeon* isolate RumEn M1 as being highly active for the incorporation of BocK. We demonstrate that genomic SCRaMbLE and high-throughput selection can tailor the genomic background of synthetic *S. cerevisiae* to improve the production of ncAA-containing protein. By pooling whole-genome sequencing data across fourteen SCRaMbLEd isolates with improved production of ncAA-containing proteins, we discovered an association of ribosomal proteins with determining genetic code expansion activity.

The exact function of many RPs remains unknown, making the causal connection of individual RPs to genetic code expansion elusive. In *S. cerevisiae*, the 80S ribosome is formed with 79 RPs that bind across the large and small subunits where they are thought to aid in rRNA folding and ribosome assembly^29^. However, there is a growing body of evidence that suggests RPs also have an integral role in modulating ribosomal properties like mRNA selectivity^30^, elongation rate^31^, and stop codon readthrough^32^. This is likely because, although the catalytic activity of the ribosome is ribozymatic^33,34^, ribosome structure and translation factor interaction may be guided by the network of ribosomal proteins^35^.

Here, we find that individual deletion of *RPS4A*, *RPS6B*, and *RPL19A* contribute to the increased production of protein containing BocK. Considering RPs are intrinsically linked to ribosome biogenesis, we hypothesize that deletion of the genes encoding these proteins lead to reduction in the pool of 80S ribosomes and that this reduced translational capacity better matches the ability of the OTS to suppress amber sites. By equilibrating the rate of translation to the rate of OTS-mediated stop codon suppression, fewer ribosome stalling events or competitive release factor terminations are incurred in which a ribosome meets an amber site that is not quickly supplied with an ncAA-tRNA. For the combined deletion of *RPL2A* and *RPL29*, the proximity of both proteins to the peptidyl transferase center (PTC) could signify an alternate biological mechanism. We assume their combined deletion produces structural modifications to the PTC that may promote ncAA translation. It is plausible that this structural effect can be specific for ncAA-tRNA pairings. We observed an improvement with BocK-RumEnPylT and PcK-RumEnPylT but not with the structurally dissimilar OMeY-*Ec*LeuT pairing. Modifying the PTC to favor ncAA-tRNA through the RP network would be an attractive method to aid genetic code expansion in yeast and higher eukaryotes, given the current inability to directly modify rRNA sequences in this domain.

For the deletion of *TMA22*, we hypothesize that the increased production of ncAA-containing protein is aided by inhibited degradation of stop codon-containing protein transcripts. Tma22p and its partner, Tma20p, bind to the 40S subunit during ribosome disassembly, preventing re-association of the 60S large subunit^36^. Removing this safeguard has been shown to disrupt the non-sense mediated decay (NMD) mRNA surveillance pathway, which targets transcripts with premature stop codons for degradation^37^. Depletion of transcripts encoding ncAAs at stop codons is an inherent concern for UAG-directed genetic code expansion given the multiple regulatory pathways that have evolved to combat non-sense mutation^38^. Previous studies have shown that deletion of an essential gene in NMD, *UPF1*, leads to significant improvements in ncAA translation by maintaining transcript abundance^39^. We did not observe improvements within the fluorescence assay for this gene deletion, however, NMD is known to be sensitive to stop codon location within the transcript.

Progress in optimizing yeast towards the production of ncAA-containing proteins is important for the transition of genetic code expansion towards industrial applications. Yeast bridge the gap between fast-growing bacterial cultures, and slow-growing mammalian cell cultures required for the production of proteins with extensive post-translational modifications^40,41^. Therefore, yeast will be important for the production of many industrially relevant proteins such as eukaryotic-derived bio-therapeutics and biocatalysts. Our SCRaMbLE pipeline has allowed the evolution of synthetic yeast strains with improved capacity to produce BocK-containing protein. By identifying the causal genomic alterations supporting improved BocK-translation, we were able to engineer from a wild-type background high-fitness yeast that were adapted towards translating BocK as well as functionally useful PcK and OMeY amino acids. However, our work highlights that these adaptations can be ncAA and OTS specific. Therefore, we anticipate that efforts to optimize of strains to produce specific ncAA-containing proteins will benefit from applying our pipeline and deconvolution strategies.

The future of genomic optimization for genetic code expansion will require continued advancements in our understanding of translation. Given the complexity of this intricate cellular process, we envisage that comprehensive cellular re-factoring will be required before orthogonal translation systems can begin to match endogenous components in translation efficiency. As highlighted in our findings, the ribosome will be a key target for continued improvement of ncAA-incorporation. By showing that the available set of RP genes can modify ribosome characteristics to increase the production of ncAA-containing proteins, we hope to open up a new line of research for the optimization of organisms toward expanded genetic codes.

## 3 Materials and Methods

### 3.1 Materials

N(*ɛ*)-Boc-L-lysine was supplied by Thermo Fisher Scientific (catalog number B21738.06), N^6^-((prop-2-yn-1-yloxy)carbonyl)-L-lysine by Activate Scientific (Cat. AS87682) and O-methyl-L-tyrosine by Thermo Fisher Scientific (Cat. 169750050). Amino acids were dissolved in Milli-Q^®^ water to give 50 mM stock solutions. For BocK and OMeY this was aided by the addition of NaOH to a final concentration of 25 mM and 50 mM respectively. Stock solutions were sterilized by filtering through a 0.2 µm PES syringe filter. *β*-estradiol was supplied by Sigma-Aldrich (Cat. E8875) and dissolved in 100% ethanol to form a 1 mM stock solution.

### 3.2 DNA Constructs

The *URA3* gene with native promoter and terminator was cloned from the pRS416 shuttle vector into pRS413 by Gibson assembly to form pRS413-URA3 (YCe6644). Codons of residues predicted to be suitable for ncAA substitution (*Fig. S1*) were mutated to the TAG triplet by Gibson assembly to form pRS413-URA3-2TAG (YCe6646) and pRS413-URA3-3TAG (YCe8031). Pyrrolysyl aminoacyl-tRNA synthetase sequences were codon optimized for expression in *S. cerevisiae* and cloned into the pRS425 shuttle vector under the control of the ADH1 promoter and terminator. Into the same plasmid, pyrrolysyl tRNA sequences were sub-cloned under expression of the *Sc*ArgT-*Sc*AspT dicistron as *Sc*ArgT-PylT^42^. All gene fragments were synthesized by Twist Biosciences. The pRS413-BXG fluorescence assay (YCe7336) was assembled by sub-cloning the fluorescent gene cassette from pCTCON2-BXG^17^ into the pRS413 shuttle vector. pCTCON2-BXG was a gift from James Van Deventer (Addgene plasmid #158127). For SCRaMbLE applications, the RumEnPylRS-RumEnPylT cassette was sub-cloned from pRS425-RumEnPylRS-RumEnPylT onto the pRS327 shuttle vector (YCe7387). Cre-EBD was cloned from pSCW11-Cre-EBD^43^ to be expressed under the PDI1 promoter to form pPDI-Cre-EBD-LEU2 (YCe7503).

### 3.3 Phylogenetic Analysis of Pyrrolysyl Homolog tRNA

tRNA sequences were aligned using the MUSCLE algorithm in MEGA-X version 11^44^. From this alignment, a maximum likelihood tree was made with the Tamura-Nei model and 1000 bootstraps.

### 3.4 Yeast Spot Assays

Strains transformed with the *URA3* growth assay were picked from single colonies and grown overnight in 10 mL selective media at 30 *^◦^*C with shaking (220 rpm). Overnight cultures were diluted to OD_600_ of 0.1, 0.01, 0.001 and 0.0001 in uracil-deficient synthetic complete media. Dilutions were spotted in 10 µL volumes onto four selective environments. These conditions were: -Ura, ncAA absent; +Ura, ncAA absent; -Ura, ncAA present; +Ura, ncAA present. Plates were incubated at 30 *^◦^*C. A picture of colony growth was taken after three and six days of incubation with a PhenoBooth^™^ from Singer Instruments.

### 3.5 Fluorescence Assay Characterization

BY4742 (MAT*α his3*Δ*1 leu2*Δ*0 lys2*Δ*0 ura3*Δ*0*) was transformed with an orthogonal translation system (pRS425-PylRS-PylT) and pRS413-BXG. As a background control, one strain was transformed with the backbone plasmids of both the orthogonal translation system and reporter assay. Single yeast colonies were taken from transformation plates in triplicate and grown to saturation over two days in 2.2 mL 96-well plates in 1 mL SC-His-Leu with glucose at 30 *^◦^*C with shaking (850 rpm). Cultures were then back-diluted to 0.1 OD_600_ in 2.2 mL 96-well plates in 1 mL SC-His-Leu with glucose and grown to mid-log phase (0.4-0.7 OD_600_). Cells were then spun down at 4000 g for five minutes and re-suspended in 50 µL sterilized Milli-Q^®^ water. These cells were then used to inoculate 1 mL of SC-His-Leu with galactose and SC-His-Leu + ncAA with galactose to 0.1 OD_600_ in a 2.2 mL 96-well plate. These cultures were incubated for 24 hours at 30 *^◦^*C with shaking (850 rpm). After 24 hours, the plate was re-suspended at 1500 rpm with an Eppendorf MixMate^®^ mixer and 200 µL was transferred to a 96-well plate. Fluorescence and OD_600_ measurements were then taken with an excitation of 479 nm and emission at 520 nm in a BioTek Synergy H1 micro-plate reader. Fluorescence of GFP was normalized to OD_600_ and averaged across three biological replicates to form a mean. Error bars were plotted as ± one standard deviation from these replicates. Auto-fluorescence was calculated in the same manner for strains transformed with two backbone plasmids and plotted as a dashed line across the x-axis.

### 3.6 *LYS2* Knockout

The *LYS2* gene was knocked out from Syn6.5 to liberate the marker for plasmid selection. A CRISPR-Cas9 system^45^ was used with a sgRNA targeting the *LYS2* gene to create a double-stranded break that was repaired with 500 bp template DNA from upstream and downstream of the *LYS2* gene (*Supplementary Information*).

### 3.7 SCRaMbLE Induction

Syn6.5 *lys2* Δ was transformed with Cre (pPDI1-Cre-EBD-LEU2), RumEn orthogonal translation system (pRS327-RumEnPylRS-RumEnPylT) and URA3 growth assay (pRS413-URA3-2TAG) with selection on SC-His-Leu-Lys media. Single colonies were picked to inoculate 10 mL SC-His-Leu-Lys overnight cultures which were incubated at 30 °C with shaking (220 rpm). The following day, cultures were back-diluted to an OD_600_ of 0.2 in 25 mL SC-His-Leu-Lys with 1 nM *β*-estradiol. Cultures were grown for 24 hours at 30 °C with shaking (220 rpm). Cultures were pelleted at 4000 g for five minutes and washed in 10 mL sterile Milli-Q^®^ water. Cells were re-suspended in Milli-Q^®^ water to an OD_600_ of 0.1. 200 µL of this culture was then plated on SC-His-Lys-Ura and SC-His-Lys-Ura +1 mM BocK Petri dishes. Plates were incubated at 30 °C for three to five days until colonies formed.

### 3.8 SCRaMbLE Selection Pipeline

Colonies from SC-His-Lys-Ura +1 mM BocK SCRaMbLE plates were re-arrayed in 96-well format with the PIXL^®^ automated colony picker from Singer Instruments. Colony selection was filtered by colony size and picked onto SC-His-Lys Nunc^TM^ OmniTrays^TM^. Plates were incubated overnight at 30 °C. Colonies were then inoculated into 96-well deep well plates with 1 mL SC-His-Lys and grown to saturation at 30 °C with shaking (850 rpm). After two days, 200 µL of the cultures were transferred to 96-well plates and spotted on selection media with the ROTOR^®^ from Singer Instruments in 7x7 grids. Cultures were assessed on SC-His-Lys-Ura and SC-His-Lys-Ura +1 mM BocK media. Plates were incubated at 30 °C for four to five days. The top ten performing strains that exhibited strong growth on SC-His-Lys-Ura +1 mM BocK and no promiscuous growth on SC-His-Lys-Ura were picked from original re-array plates and cultured in 10 mL SC-His-Lys for 24 hours at 30 °C with shaking (220 rpm). The strains were then spot-assayed on the four selective conditions containing either uracil/BocK.

### 3.9 Yeast Growth Rate Assay

Single colonies were picked in triplicate and inoculated into 1 mL SC-His-Lys in a 96-deep-well plate. Plates were incubated to saturation for two days at 30 °C with shaking (850 rpm). Cultures were diluted 1:100 in 200 µL media in a 96-well plate. Assay plates were cultured at 30 °C with shaking (200 rpm) for 72 hours in a BioTeK Synergy H1 micro-plate reader. Absorbance at 600 nm was measured every 10 minutes. The growth rate as doublings per hour was calculated as follows: OD_600_ values were subtracted by the background absorbance of a media-only control. A log_2_ was taken of OD_600_ measurements between the values of 0.2 and 0.6. The equation for the line of best fit as a linear least squares regression was calculated for the log_2_(OD_600_) values against time. The doubling rate was derived from gradient of this line and converted to doublings per hour by calculating 1 / rate. Data was plotted as the mean of three biological replicates ± one standard deviation.

### 3.10 Nanopore Sequencing

Genomic DNA was isolated from strains using the high-molecular weight NucleoBond kit from Macherey-Nagel (Cat. 740160.20). gDNA was sequenced with long-read sequencing using the MinION Mk1B from Oxford Nanopore Technologies. 1 µg of high-molecular weight genomic DNA from each sample was prepared for ligation sequencing using the Native Barcoding kit 24 V14 (Cat. No. SQK-NBD114.24). Barcoded gDNA was quantified with the Qubit^TM^ 4 Fluorometer with equimolar quantities of each sample being pooled into a library. The combined gDNA library was loaded onto a MinION flow cell (R10.4.1) on a MinION Mk1B device. Guppy version 6.4.6 (GPU) was used for base calling with default parameters and inclusion of de-multiplexing, adaptor and barcode trimming and read splitting.

### 3.11 Structural Variation Detection

De-mutiplexed reads for each sample were pre-processed using Chopper version 0.3.0^46^ to remove low quality (Phred *<* 9) and short (*<* 500 bp) reads. Reads were aligned to the reference genome using Minimap2 version 2.17^47^. Sequence alignment map files were sorted, indexed and compressed using SAMtools version 1.15.1^48^. Putative structural variants were called using Sniffles version 2.0.6^49^. Structural variation was visualized and assessed using Integrative Genomics Viewer version 2.16.2^50^.

### 3.12 Interactome Mapping

Direct protein-protein interactions were taken from affinity purification - mass spectrometry data^51^. Interactions were mapped with a custom script for visualization in Obsidian.md. Genes that were disrupted during SCRaMbLE were annotated on the graph and color-coded by strain. Clusters of cellular functions were assessed and labeled qualitatively.

### 3.13 Gene Set Enrichment Analysis

SCRaMbLEd genes were analyzed using the gene-ontology (GO) framework. Genes within the sets were classified into GO protein functional terms using the PANTHER classification system (version 19.0)^52^. A PANTHER statistical over-representation test compared the gene lists from the two sets against a reference list of genes found on the synthetic chromosomes of Syn6.5, excluding rRNA genes. The comparison was performed with a Fisher’s exact test and no correction for false discovery. GO-terms with a *>* 2-fold enrichment and a P-value *<* 0.05 were plotted.

### 3.14 Yeast Knockout Strains

Strains of BY4742 (MAT*α his3*Δ*1 leu2*Δ*0 lys2*Δ*0 ura3*Δ*0*) from the yeast knockout collection^53^ were recovered from glycerol stocks on YPD + 200 µg/mL G418. Strains were verified for correct gene single deletion by PCR with two sets of confirmation primers from the collection (A+B, C+D). Single colonies were re-suspended in 200 µL 20 mM NaOH and incubated at 95 °C for 15 minutes. 10 µL of this solution was used as template for the PCR reaction. PCRs were performed with the 2X DreamTaq DNA polymerase master mix from Thermo Scientific (Cat. EP0705) in a 25 µL total reaction volume. The standard DreamTaq protocol was followed with an annealing temperature of 55 °C and an extension time of 30 seconds. PCR reactions were run for 20 minutes at 100 V on a 1% agarose gel. The double knockout, *rpl2a*Δ*rpl29* Δ, was made using a NATMX cassette with 50 bp homology upstream and downstream of *RPL2A* to remove the gene from the *rpl29* Δ strain of the knockout collection. Transformants were selected on YPD + 200 µg/mL G418, + 100 µg/mL NAT.

### 3.15 msGFP Assay Characterization

Strains were transformed with pRS425-RumEnPylT-RumEnPylRS and either pRS423-pTDH3-msGFP or pRS423-pTDH3-msGFP-N149TAG assays. As a control for auto-fluorescence, BY4742 was also transformed with two empty backbones. Single colonies were picked into 700 *µ*L SC-His-Leu in a deep well plate and grown to saturation at 30 °C with shaking (850 rpm). Once saturated, 10 *µ*L was used to inoculate 990 *µ*L of SC-His-Leu with and without 1 mM BocK in a deep well plate. Plates were cultured for 24 hours at 30 °C with shaking (850 rpm) after which 200 *µ*L was transferred to a 96-well plate from which msGFP expression (ex. 479 nm, em. 520 nm) and absorbance at 600 nm was measured on a BioTek Synergy H1 micro-plate reader. Final GFP fluorescence values were calculated by normalizing msGFP expression to OD_600_ followed by subtraction of a normalized auto-fluorescence control strain.

### 3.16 msGFP Production and Purification

Strains were transformed with pRS423-msGFP-N149TAG and pRS425-RumEnPylT-PylRS plasmids. As control, BY4742 was transformed with pRS423-msGFP and pRS425-RumEnPylT-RumEnPylRS. A single colony of freshly transformed cells was cultured in 5 mL of SC-His-Leu media for 18 hours at 30 °C. 500 *µ*L of the starter culture was used to inoculate 500 mL SC-His-Leu media supplemented with 1 mM BocK, and cultures were grown at 30 °C, 200 rpm for 40 hours. The cells were harvested by centrifugation at 4000 g for 15 min. Cells were resuspended in 2.5 mL/g of pellet in Y-PER Yeast Protein Extraction Reagent supplied by Thermo Scientific (Cat. 78990) and agitated for 20 minutes at room temperature. Cell lysates were cleared by centrifugation (4000 g for 15 minutes) and the supernatants subjected to affinity chromatography using Ni-NTA agarose supplied by Qiagen (Cat. 30210). After washing (50 mM HEPES, 300 mM NaCl, pH 7.5 and 50 mM imidazole), purified protein was eluted using 50 mM HEPES, 300 mM NaCl, pH 7.5 and 250 mM imidazole. Proteins were desalted using 10DG desalting columns supplied by Bio-Rad (Cat. 7322010) with PBS pH 7.4 and analyzed by SDS-PAGE. Protein concentrations were determined by measuring the absorbance at 280 nm. Proteins were concentrated and flash-frozen in liquid nitrogen and stored at -80 °C.

### 3.17 Mass Spectrometry

Purified protein samples were diluted to a final concentration of 0.5 mg/mL in PBS pH 7.4. MS analysis was performed using a 1200 series Agilent LC system, with a 5 *µ*L injection into 5% acetonitrile (with 0.1% formic acid), and desalted inline for 1 minute. Protein was eluted over 1 minute using 95% acetonitrile with 5% water. The resulting multiply charged spectrum was analyzed using an Agilent QTOF 6510 instrument and deconvoluted using Agilent MassHunter software.

## Acknowledgments

This work is supported by UKRI Engineering Biology Mission award (BB/Y00812X/1), ERC consolidator award (UKRI Frontier Research Guarantee EP/Y023722/1) to A.P.G; UKRI Engineering Biology Transition Award (BB/W014483/1), ERC consolidator award (UKRI Frontier Research Guarantee EP/Y024753/1), EPSRC Open Plus Fellowship (EP/V05967X/1) to Y. C; and EPSRC DTG scholarship to J. E. J. S. We thank the University of Manchester Faculty of Science & Engineering Mass Spectrometry and Separations Facility (RRID SCR 024761), in particular we thank Kathleen Cain and Michael Trelore. The plasmids pCTCON2-BYG (Addgene plasmid # 158126) and pCTCON2-BXG (Addgene plasmid # 158127) were gifts from James Van Deventer.

## Contributions

Y.C., A.P.G., S.H. and J.E.J.S. conceptualized the study that Y.C., A.P.G. and S.A.H. supervised and acquired funding for. J.E.J.S performed the majority of the investigation, to which E.R.M. contributed to by performing all protein purifications, M.M. contributed to by assembling genomic data from NGS workflows, and S.H. contributed to by developing the *URA3* growth assay. All authors contributed to the development of the methodology. Formal analysis, validation and visualization was performed by J.E.J.S aided by data curation from J.E.J.S and M.M.. The original draft was written by J.E.J.S with all authors contributing to review and editing. Resources were supplied by A.P.G and Y.C., with project administration by Y.C..

## 7. Supplementary Information

### 7. 1 Supplementary Figures

**Figure S1:**
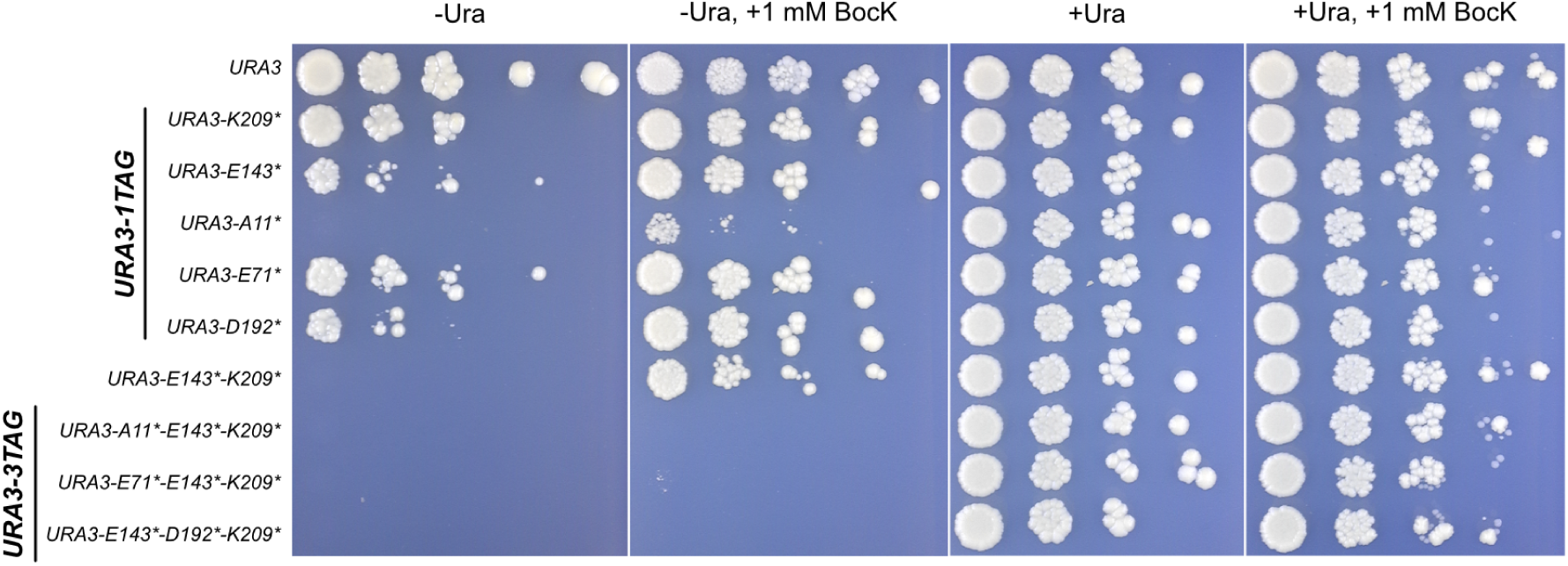
Characterization of the *URA3* Growth Assay. Sites within the Ura3p protein structure (PDB: 1DQW) were identified to be permissible for amino acid substitution. These positions were assessed *in vivo* by replacing the corresponding codon for the TAG stop codon to form *URA3-1TAG* assays. To increase selection pressure, single sites were stacked to form *URA3-2TAG* and *URA3-3TAG* assays. Strains of BY4742 were transformed with the growth assays and pRS327-RumEn-RumEnPylT. Transformants were cultured in SC-His-Lys media and spotted on SC-His-Lys in the presence and/or absence of uracil/1 mM BocK. Plates were cultured at 30 °C for five days.

**Figure S2:**
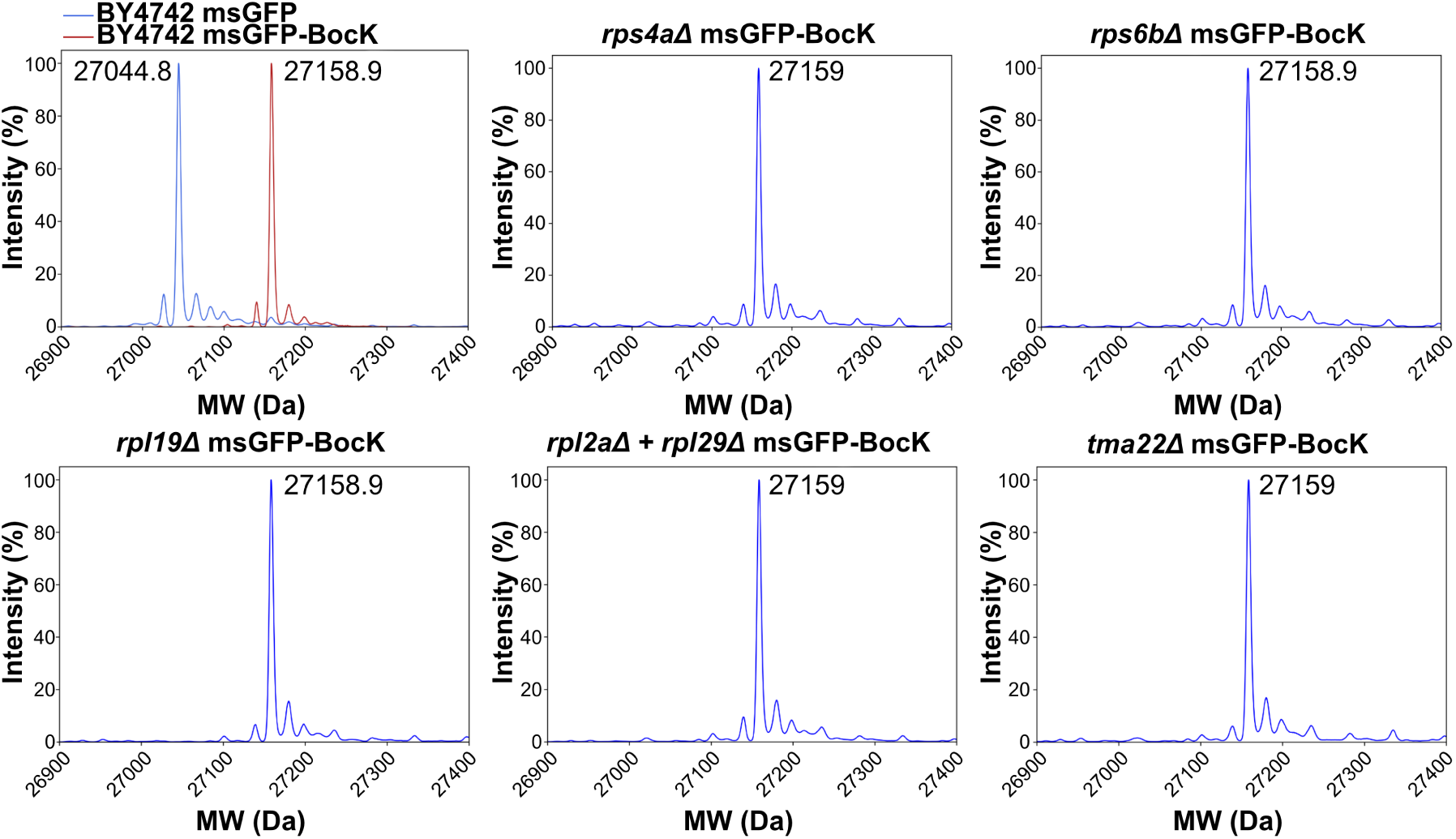
Mass Spectra of msGFP Protein. Mass spectra of msGFP-BocK isolated from wild-type BY4742 and knockout strains transformed with pRS423-msGFP-N149TAG and pRS425-RumEnPylT-RumEnPylRS. As control full length ms-GFP was purified from the wild-type BY4742 strain transformed with pRS423-msGFP + pRS425-RumEnPylT-RumEnPylRS.

### 7.2 Pyrrolysyl Synthetase Amino Acid Sequences

Organism = Methanogenic archaeon ISO4-G1, Protein = G1PylRS

MVVKFTDSQIQHLMEYGDNDWSEAEFEDAAARDKEFSSQFSKLKSANDKGLKDVIANPRN DLTDLENKIREKLAARGFIEVHTPIFVSKSALAKMTITEDHPLFKQVFWIDDKRALRPMHA MNLYKVMRELRDHTKGPVKIFEIGSCFRKESKSSTHLEEFTMLNLVEMGPDGDPMEHLKM YIGDIMDAVGVEYTTSREESDVYVETLDVEINGTEVASGAVGPHKLDPAHDVHEPWAGIGF GLERLLMLKNGKSNARKTGKSITYLNGYKLD

Organism = *Methanomassiliicoccus luminyensis* 1, Protein = *Lum1* PylRS

MTRLTPAQAQRIREMGGTVDPSLAFSSEAERESAFQRISADLQGANLAKIRRCAEAPERHPI GSLENTLACALAAKGFIEVKTPMMIPADGLVKMGIDESHPLWNQVFWVGPKKALRPMLAP NLYFLMRHLRRSVPAPLLLFEIGPCFRKESRGSNHLEEFTMLNLVELAPQADATERLKEHIA TVMNAVGLPYELVVEGSEVYGTTIDVEVDGVELASGAVGPLPMDKPHGITEPWAGVGFGL ERIALMRTKEQNIKKVGRSLVYVNGARIDI

Organism = *Methanoplasma termitum*, Protein = *Term*PylRS

MSIGFTPSQIQKLREFGEDPRDNSTYQNVEQRDKAFSKLMSDLVSSNEKEIAGMLRSPSRHQ LAALEEDLAAALIARGFIEVKTPAFVSVASLEKMTITPEHPLYKQVFMIDDKRCLRPMHAM NLYYVMRKLRDHTDGPVKIFEIGSCFRKESHSGSHLEEFTMLNLVELGPEGDATEALKDHIG AVMKVTGLEYTLVREESDVYVETLDVEIDGGEVASGAVGPHVLDNAHDIHEPWSGIGFGLE RLLMIMNEKSTVKKTGRSLSYLNGAKIN

Organism = Methanogenic archaeon ISO4-H5, Protein = H5PylRS

MTCKLTDPQIQRLREYGHEPKNESEFETEEERDKAFTKMMSKLQRENEKGIRDMIANPRH HRLMELELQLSEALIKEGFIEVKTPILISKAELAKMTIDENHPLYQQVFWVDDKRCLRPMHA INLYNIMRELRGHTDGPVKFFEIGSCFRAESHSNDHLEEFTMLNLVDMGPQGDTTEKIKHYI DIVMKTIGLDYELVHEESDVYKETIDVEVDGEEVCSAAVGPHYLDKAHNINEPWCGAGFGL ERLIMMRDGDGSVKKTGKSVNYLNGYKIN

Organism = *Methanomassiliicoccales archaeon* PtaU1.Bin030, Protein = 030PylRS

MVIEWSPSQKQRLRELGRADEGGMEFETVVERDEAFTKEVAYYQSINRKEIRNIQERRERHLLAKVEENIAEALIADGFLEVRTPTIISGNALVKMGIDHNHPLREQVFWLDGSRCLRPMLAPNLYFLMRHLKRNVRMPLQMFEIGTCYRKESHGSNHLEEFTMLNLVEMASMDDPAVRLRHHIQTVMGAIGLEYELSECESDVYGRTIDVEVNGVEVASAALGPHKLDPAHGITDAWSGVGFGLERLLMVKNAENNIKKVGRSLIYLGGARLDI

Organism = *Methanomethylophilus alvus*, Protein = *Alv* PylRS

MTVKYTDAQIQRLREYGNGTYEQKVFEDLASRDAAFSKEMSVASTDNEKKIKGMIANPSRHGLTQLMNDIADALVAEGFIEVRTPIFISKDALARMTITEDKPLFKQVFWIDEKRALRPMLAPNLYSVMRDLRDHTDGPVKIFEMGSCFRKESHSGMHLEEFTMLNLVDMGPRGDATEVLKNYISVVMKAAGLPDYDLVQEESDVYKETIDVEINGQEVCSAAVGPHYLDAAHDVHEPWSGAGFGLERLLTIREKYSTVKKGGASISYLNGAKIN

Organism = *Candidatus Methanomethylophilus* sp. 1R26, Protein = 1R26PylRS

MAEHFTDAQIQRLREYGNGTYKDMEFADVSAREKAFTKLMSDASRDNESALKGMIAHPARQGLSRLMNDIADALVADGFIEVRTPIIISKDALAKMTITPDKPLFKQVFWIDDKRALRPMLAPSLYTVMRSLRDHTDGPVKIFEMGSCFRKESHSGMHLEEFTMLNLVDMGPAGDATESLKKYIGIVMKAAGLPDYQLVHEESDVYKETIDVEINGQEVCSAAVGPHYLDAAHDVHEPWAGAGFGLERLLTIRQGYSTVMKGGASTTYLNGAKMD

Organism = *Methanomassiliicoccales archaeon* RumEn M1, Protein = RumEnPylRS

MTIEWTPSQKQRLKELGIDSDQDYTINNIQEREEVFSRLVTRRQSEGRRAIRSMMEHPVRHKLAQLEQDLAQALVDDGFLEFRTPTIITRSALEKMGIGREHPLHEQVFWLDEKRCLRPMLAPNLYYVMRHLKRNAKGPVKLFEIGTCYRKESHGSNHLEEFTMLNLVELDPAGDAREQLRKHISTIMNTIGLDYELVSCSSDVYVETTDVEVNGVEVASGAIGPHKLDPAHGIKAPWAGVGFGLERLLMLKHGEDNVKKVGRSLIYLQGVRLDI

Organism = *Methermicoccus shengliensis*, Protein = *Sheng* PylRS

MIGFTDTQVQRLKELGGDQKIVGCRFSSVADRDEVFETTVKNLVEENREKLRRMAHSPSRCSLFELEDRLASTLVGMGFMEVATPMLLSASNLKKMGIDESHPLWEQVFWVDKKRCMRPMLAPNLYFLLKHLKRNIKKPVRIFEIGPCFRKESQGSRHLEEFTMLNLVELAPDCEPTERLTELIEKVMGQIGLEYRLKNESSEVYGNTVDVEVDGVEVASGAVGPHFLDGAYGITDAWVGVGFGLERLLMVMEGHSNIRKVGRSLVYLNGARIDI

Organism = *Methanomassiliicoccus intestinalis*, Protein = *Int* PylRS

MPVEWTASQKQRLKELGIPAEADRIFNDTKEREEVFKDITSEHLSKVRKDIKHMLDYPERHQLSQIESILAQALVDNGFIEVKTPSIISRSALEKMGIDRSHPLHEQVFWLDEKRCLRPMLAPNLYFMMRHMYRYSKGPLRLFEIGSCFRKESKGSNHLEEFTMLNLVEMAPDNDPADQLLVHIKTIMDALGLEYSLVECESDVYVKTLDVEIDGVEVASGAVGPHKLDPAHGITQSWAGVGFGLERLSMMKYGMDNIKKSGRSLIYLRGVRLDI

Organism = *Methanomassiliicoccus luminyensis* 2, Protein = *Lum2* PylRS

MVRDRQASTATAAGFDSRALRSQHYESEDGGWQLIFDMTPSQKQRLRELGRVPDEGAAFSTAEDRDAAFIKEVAYYQSYNRNVVRDALDAPKRHPLSHMEEVLAQALVDEGFLDVKTPTIISGDSIRKMGISCAHPLNKQIFWVDGTRCLRPMLAPNLYFLMRHLKRNAQLPLRLFEIGPCYRIETHGSDHLEEFTMLNLVELAPQGDPLAQLHHHIATVMGAVGLDYQLCECDSEVYSRTIDVEVDGSEVASAALGPHALDRAHGIEDPWVGVGFGLERLLMSKSAESNIRKVGRSLIYLQGA RIDV

### 7.3 Pyrrolysyl tRNA Sequences

*Sc*ArgT-*Sc*PylT Dicistron Expression System Capitalized = *Sc*ArgT, N = PylT

accggtatcgatgtgtgttatatgtacctctgctttgcagtataagaaatttacatttatttctgactaataacaccttggtgccccaacggtaaacaacttgtatcagttctcataagtgcggccattttatgcaatacaggctgcattatttcaccagccgtgaaaatccgaaaattgtagtaattgaaagcgtaattaggttttactataataaagtagtaaaaccttcaacaaatagtaGCTCGCGTGGCGTAATGGCAACGCGTCTGACTTCTAATCAGAAGATTATGGGTTCGACCCCCATCGTGAGTGctttgtttctNNNNNNNNNNNNNNNNNNNNNNNNNattttttggctactcctgtagttattcttcattaatgctttgttaacgctag

Organism = Methanogenic archaeon ISO4-G1, Gene = G1PylT

GGAGGGCGCTCCGGCGAGCAAACGGGTCTCTAAAACCTGTAAGCGGGGTTCGACCCC CCGGCCTTTCG

Organism = *Methanomassiliicoccus luminyensis* 1, Gene = *Lum1* PylT

GGAGTGTTGGTTCGGCGACCACCAGGCCTCTACAGCCACGGCAGCCGGGTTCGACTCC CGGGCACTTCG

Organism = *Methanoplasma termitum*, Gene = *Term*PylT

GGAGGGCGCTCCGGCGAGCAAACGGGTCTCTAAAACCTGTAAGCGGGGTTCGACCCC CCGGCCTTTCG

Organism = Methanogenic archaeon ISO4-H5, Gene = H5PylT

GGAGGGCGCTCCGGCGAGCAAACGGGTCTCTAAAACCTGTAAGCGGGGTTCGACCCC CCGGCCTTTCG

Organism = *Methanomassiliicoccales archaeon* PtaU1.Bin030, Gene = 030PylT

GGAGGGTTGGTCCGGGACCACCTGGCCTCTACAGCTAAGGCAGCCGGGTTCAACTCCC GGGCCCTTCG

Organism = *Methanomethylophilus alvus*, Gene = *Alv* PylT

GGGGGACGGTCCGGCGACCAGCGGGTCTCTAAAACCTAGCCAGCGGGGTTCGACGCC CCGGTCTCTCG

Organism = *Candidatus Methanomethylophilus* sp. 1R26, Gene = 1R26PylT

GGGGGACGATCCGGCGATCAGCGGGTCTCTAAAACCTAGCCAGCGGGGATCGACACC CCGGTCTCTCG

Organism = *Methanomassiliicoccales archaeon* RumEn M1, Gene = RumEnPylT

GGAGTGTTGGTCCGGAGACCACCAGGCCTCTACAGCCGCGGCAGCCGGGTTCGACTC CCGGGCACTTCG

Organism = *Methermicoccus shengliensis*, Gene = *Sheng* PylT

GGAGGGTTGGTCCGGGACCGCCAGGCCTCTACAGCCACGGTAGCTGGGTTCGACTCCCAGGCCCTTCG

Organism = *Methanomassiliicoccus intestinalis*, Gene = *Int* PylT

GGAGTGTTGGTCCGGGACCACCAGGCCTCTACAGCCACGGCAGCCGGGTTCAACTCCCGGGCACTTCG

Organism = *Methanomassiliicoccus luminyensis* 2, Gene = *Lum2* PylT

GGAGGGTTGGTCAGGGACCGCCAGGCCTCTACAGCCACGGCAGCCGGGTTCGACTCCCGGGCCCTTCG

### 7.4 *URA3* Growth Assay

*URA3-2TAG* Assay = *URA3-E143*-K209\**

*URA3-3TAG* Assay = *URA3-E71*-E143*-K209\**

Ura3 Protein Sequence

MSKATYKERAATHPSPVAAKLFNIMHEKQTNLCASLDVRTTKELLELVALGPKICLLKTHV DILTDFSM<u>E</u>GTVKPLKALSAKYNFLLFEDRKFADIGNTVKLQYSAGVYRIAEWADITNAHG VVGPGIVSGLKQAAEEVTK<u>E</u>PRGLLMLAELSCKGSLSTGEYTKGTVDIAKSDKDFVIGFIAQ RDMGGRDEGYDWLIMTPGVGLDD<u>K</u>GDALGQQYRTVDDVVSTGSDIIIVGRGLFAKGRDA KVEGERYRKAGWEAYLRRCGQQN

### 7.5 msGFP Fluorescence Assay

TDH3 Promoter

CGAGTTTATCATTATCAATACTGCCATTTCAAAGAATACGTAAATAATTAATAGTAGTGATTTTCCTAACTTTATTTAGTCAAAAAATTAGCCTTTTAATTCTGCTGTAACCCGTACATGCCCAAAATAGGGGGCGGGTTACACAGAATATATAACATCGTAGGTGTCTGGGTGAACAGTTTATTCCTGGCATCCACTAAATATAATGGAGCCCGCTTTTTAAGCTGGCATCCAGAAAAAAAAAGAATCCCAGCACCAAAATATTGTTTTCTTCACCAACCATCAGTTCATAGGTCCATTCTCTTAGCGCAACTACAGAGAACAGGGGCACAAACAGGCAAAAAACGGGCACAACCTCAATGGAGTGATGCAACCTGCCTGGAGTAAATGATGACACAAGGCAATTGACCCACGCATGTATCTATCTCATTTTCTTACACCTTCTATTACCTTCTGCTCTCTCTGATTTGGAAAAAGCTGAAAAAAAAGGTTGAAACCAGTTCCCTGAAATTATTCCCCTACTTGACTAATAAGTATATAAAGACGGTAGGTATTGATTGTAATTCTGTAAATCTATTTCTTAAACTTCTTAAATTCTACTTTTATAGTTAGTCTTTTTTTTAGTTTTAAAACACCAAGAACTTAGTTTCGAATAAACACACATAAACAAACAAAG

msGFP Protein Sequence

MDSTESLFTGVVPILVELDGDVNGHKFSVRGEGEGDATNGKLTLKFICTTGKLPVPWPTLVTTLTYGVQCFSRYPDHMKQHDFFKSATPEGYVQERTITFKDDGTYKTRAEVKFEGDTLVN RIELKGIDFKEDGNILGHKLEYNYNSH<u>N</u>VYITADKQKNGIKANFKIRHNVEDGSVQLADHYQ QNTPIGDGPVLLPDNHYLSTQSKLSKDPNEKRDHMVLLEFVTAAGITGGSGGSHHHHHH

KRR1 Terminator Sequence

TTTACACCGCTACGGAATCCTTCATATATGTCTATATATATATAGAAACACACACACATTTCGTTTACTTAGATATGTTTGTAAAGTGAAAGAAATTCTTACTAGGACACCAACTCTACTGGCTCTCATATGCAGAGAGATATACATCAATACTTGAAATTTTCCTTATCCTTCTTCGACGCGTCATGAAGAGATAGAAT

### 7.6 PDI1-Cre-EBD Expression System

PDI1 Promoter + 5’ UTR Sequence

GCCCCCTGAAAGGGGTTGACCGTCCGTCGGCACACCACTTATAATGCGGGGTGCAAGCGCCGCGTCTAAAATTTTTTTTTTTTCCATTTTTGTCGTTATTGTTATTTCCCGTTTTTTGTTTTTTTTGATTTTTTCGGAGCGACAAACCTTTCGAAACACGTGTCCTGAAAATTATCCTGGGCTGCACGTGATAATATGTTACCCTGTCGGGCGGCGCCTCTTTTTCCCTTTTCTCTCACTAGTCTCTTTTTCCAATTTGCCACCGTGTAGCATTTTGTTGTGCTGTTACAACCACAACAAAACGAAAAACCCGTATGGACATACATATATATATATATATATATATATATTTTGTTACGCGTGCATTTTCTTGTTGCAAGCAGCATGTCTAATTGGTAATTTTAAAGCTGCCAAGCTCTACATAAAGAAAAACATACATCTATCCCGTTGATGCGAGACGACTGACCCATCATCCTGTACACTTTACTTAAAACCATTATCTGAGTGTTA

Cre Protein Sequence

MSNLLTVHQNLPALPVDATSDEVRKNLMDMFRDRQAFSEHTWKMLLSVCRSWAAWCKLNNRKWFPAEPEDVRDYLLYLQARGLAVKTIQQHLGQLNMLHRRSGLPRPSDSNAVSLVMRRIRKENVDAGERAKQALAFERTDFDQVRSLMENSDRCQDIRNLAFLGIAYNTLLRIAEIARIRVKDISRTDGGRMLIHIGRTKTLVSTAGVEKALSLGVTKLVERWISVSGVADDPNNYLFCRVRKNGVAAPSATSQLSTRALEGIFEAAHRLIYGAEDDSGQRYLAWSGHSARVGAARDMARAGVSIPEIMQAGGWTNVNIVMNYIRNLDSETGAMVVLLEDGD

*β*-estradiol Binding Domain (EBD) Protein Sequence

LEPSAGDMRAANLWPSPLMIKRSKENSLALSLTADQMVSALLDAEPPILYSEYDPTRPFSEASMMGLLTNLADRELVHMINWAKRVPGFVDLTLHDQVHLLECAWLEILMIGLVWRSMEHPGKLLFAPNLLLDRNQGKCVEGMVEIFDMLLVTSSRFRMMNLQGEEFVCLKSIILLNSGVYTFLSSTLKSLEEKDHIHRVLDKITDTLIHLMAKAGLTLQQQHQRLAQLLLLLSHIRHMSNEGMEHLYSMKCKNVVPLYDLLLEMLDAHRLHAPTSRGGASVEETDQSHLATAGSTSSHSLQKYYITGEAEGFPATV

Terminator Sequence

GTAGAATGCCTGATATTGACTCAATATCCGTTGCGTTTCCTGTCAAAAGTATGCGTAGTGCTGAACATTTCGTGATGAATGCCACCGAGGAAGAAGCACGGCGCGGTTTTGCTAAAGTGATGTCTGAGTTTGGCGAACTCTTGGGTAAGGTTGGAATTGTCGACCTCGAGTCATGTAATTAGTTATGTCACGCTTACATTCACGCCCTCCCCCCACATCCGCTCTAACCGAAAAGGAAGGAGTTAGACAACCTGAAGTCTAGGTCCCTATTTATTTTTTTATAGTTATGTTAGTATTAAGAACGTTATTTATATTTCAAATTTTTCTTTTTTTTCTGTACAGACGCGTGTACGCATGTAACATTATACTGAAAACCTTGCTTGAGAAGGTTTTGGGACGCTCG

### 7.7 *LYS2* Knockout

*LYS2* gDNA Target Sequence

GGCAACGGTTCATCATCTCA

*LYS2* Upstream Donor Homology

AGCAGTTGCTTTCTCCTATGGGAAGAGCTTTCTAAGTCTGAAGAAGTAAACAGTTCTTTGCTATTTCACACTTCCTGGTTGATGGTCACTTGCTGCCTGAAATATATATATATGTATGACATATGTACTTGTTTTCTTTTTTGTGCCTTTGTTACGTCTATATTCATTGAAACTGATTATTCGATTTTCTTCTTGCTGACCGCTTCTAGAGGCATCGCACAGTTTTAGCGAGGAAAACTCTTCAATAGTTTTGCCAGCGGAATTCCACTTGCAATTACATAAAAAATTCCGGCGGTTTTTCGCGTGTGACTCAATGTCGAAATACCTGCCTAATGAACATGAACATCGCCCAAATGTATTTGAAGACCCGCTGGGAGAAGTTCAAGATATATAAGTAACAAGCAGCCAATAGTATAAAAAAAAATCTGAGTTTATTACCTTTCCTGGAATTTCAGTGAAAAACTGCTAATTATAGAGAGATATCACAGAGTTACTCACTA

*LYS2* Downstream Donor Homology

GGTTGAGCATTACGTATGATATGTCCATGTACAATAATTAAATATGAATTAGGAGAAAGACTTAGCTTCTTTTCGGGTGATGTCACTTAAAAACTCCGAGAATAATATATAATAAGAGAATAAAATATTAGTTATTGAATAAGAACTGTAAATCAGCTGGCGTTAGTCTGCTAATGGCAGCTTCATCTTGGTTTATTGTAGCATGAATCATATTTGCCTTTTTTTCCTGTAATTCAATGATTCTTGCTTCTATACTATCCTCAATGCAAAACCTTGTGATCTTCACAGGTCGATACTGACCAATTCTATGAACTCTATCACCACTTTGCCATTCAACACTAGGGTTCCACCATGGGTCTAAAATGAATACTTGCGAAGCTTCACAAAGATTCAAAGCAACACCGCCCGCCTTTAAACTGACCAAGAAAACCTCGCATTGAATGTTGTTCATGAAATACTTGATGGTTTCATCTCTTTGCGTCGGTGACATACTACCCTGA

**Table S1:** Strain IDs of SCRaMbLE-Related Strains.

| Strain | ID | Genotype |
| --- | --- | --- |
| Syn6.5 | YCy5385 | *MATα ura3Δ0 leu2Δ0 his3Δ1 lys2Δ*<br>ΔsynII::tS(AGA)BΔ0::tRNA-array<br>synIII::tP(AGG)CΔ0::tRNA-array synIII.WT-*SUP61*<br>ho::syn*SUP61*-HSX1 synV.tK(CUU)E1Δ::tRNA-array<br>synVI.tA(AGC)FΔ::tRNA-array synIXR<br>synX.tL(UAA)JΔ0::tRNA-array<br>synXII.tR(CCG)LΔ::XII-tRNA-array |
| D7 | YCy5406 | scrSyn6.5 *MATα ura3Δ0 leu2Δ0 his3Δ1 lys2Δ* |
| 4D3 | YCy5669 | scrSyn6.5 *MATα ura3Δ0 leu2Δ0 his3Δ1 lys2Δ* |
| 7A3 | YCy5672 | scrSyn6.5 *MATα ura3Δ0 leu2Δ0 his3Δ1 lys2Δ* |
| 7C8 | YCy5673 | scrSyn6.5 *MATα ura3Δ0 leu2Δ0 his3Δ1 lys2Δ* |
| 7E5 | YCy5674 | scrSyn6.5 *MATα ura3Δ0 leu2Δ0 his3Δ1 lys2Δ* |
| 9D8 | YCy5676 | scrSyn6.5 *MATα ura3Δ0 leu2Δ0 his3Δ1 lys2Δ* |
| 1D5 | YCy5683 | scrSyn6.5 *MATα ura3Δ0 leu2Δ0 his3Δ1 lys2Δ* |
| 6B12 | YCy5686 | scrSyn6.5 *MATα ura3Δ0 leu2Δ0 his3Δ1 lys2Δ* |
| 8B12 | YCy5687 | scrSyn6.5 *MATα ura3Δ0 leu2Δ0 his3Δ1 lys2Δ* |
| 5H3 | YCy5719 | scr7A3 *MATα ura3Δ0 leu2Δ0 his3Δ1 lys2Δ* |
| 8B5 | YCy5723 | scr7A3 *MATα ura3Δ0 leu2Δ0 his3Δ1 lys2Δ* |
| 4C2 | YCy5776 | scrD7 *MATα ura3Δ0 leu2Δ0 his3Δ1 lys2Δ* |
| 7A12 | YCy5783 | scrD7 *MATα ura3Δ0 leu2Δ0 his3Δ1 lys2Δ* |
| 8A2 | YCy5785 | scrD7 *MATα ura3Δ0 leu2Δ0 his3Δ1 lys2Δ* |
| 4F1 | YCy5888 | scrD7 *MATα ura3Δ0 leu2Δ0 his3Δ1 lys2Δ* |
| BY4742 *rps4aΔ* | YCy6392 | *MATα ura3Δ0 leu2Δ0 his3Δ1 lys2Δ rps4aΔ* |
| BY4742 *rps6bΔ* | YCy6391 | *MATα ura3Δ0 leu2Δ0 his3Δ1 lys2Δ rps6bΔ* |
| BY4742 *rpl2aΔ* | YCy6789 | *MATα ura3Δ0 leu2Δ0 his3Δ1 lys2Δ rpl2aΔ* |
| BY4742 *rpl19aΔ* | YCy6687 | *MATα ura3Δ0 leu2Δ0 his3Δ1 lys2Δ rpl19aΔ* |
| BY4742 *rpl29Δ* | YCy6790 | *MATα ura3Δ0 leu2Δ0 his3Δ1 lys2Δ rpl29Δ* |
| BY4742 *rpl2aΔ rpl29Δ* | YCy6649 | *MATα ura3Δ0 leu2Δ0 his3Δ1 lys2Δ rpl2aΔ rpl29Δ* |
| BY4742 *tma22Δ* | YCy6103 | *MATα ura3Δ0 leu2Δ0 his3Δ1 lys2Δ tma22Δ* |

